# Reduced Neutralization of SARS-CoV-2 Omicron Variant in Sera from SARS-CoV-1 Survivors after 3-dose of Vaccination

**DOI:** 10.1101/2022.05.21.492903

**Authors:** Xuesen Zhao, Danying Chen, Xiaohua Hao, Yaruo Qiu, Juan Du, Yuanyuan Zhang, Fan Xiao, Xinglin Li, Yanjun Song, Rui Song, Xi Wang, Ronghua Jin

**Affiliations:** Beijing Key Laboratory of Emerging Infectious Diseases, Institute of Infectious Diseases, Beijing Ditan Hospital, Capital Medical University, Beijing 100015, China; Beijing Institute of Infectious Diseases, Beijing, 100015, China; National Center for Infectious Diseases, Beijing Ditan Hospital, Capital Medical University, Beijing 100015, P.R. China

## Abstract

Recent studies found that Omicron variant escapes vaccine-elicited immunity. Interestingly, potent cross-clade pan-sarbecovirus neutralizing antibodies were found in survivors of the infection by SARS-CoV-1 after BNT162b2 mRNA vaccination (***N Engl J Med. 2021 Oct 7;385(15):1401-1406***). These pan-sarbecovirus neutralizing antibodies were observed to efficiently neutralize the infection driven by the S protein from both SARS-CoV and multiple SARS-CoV-2 variants of concern (VOC) including B.1.1.7 (Alpha), B.1.351 (Beta), and B.1.617.2 (Delta). However, whether these cross-reactive antibodies could neutralize the Omicron variant is still unknown. Based on the data collected from a cohort of SARS-CoV-1 survivors received 3-dose of immunization, our studies reported herein showed that a high level of neutralizing antibodies against both SARS-CoV-1 and SARS-CoV-2 were elicited by a 3rd-dose of booster vaccination of protein subunit vaccine ZF2001. However, a dramatically reduced neutralization of SARS-CoV-2 Omicron Variant (B.1.1.529) is observed in sera from these SARS-CoV-1 survivors received 3-dose of Vaccination. Our results indicates that the rapid development of pan-variant adapted vaccines is warranted.

## To the Editor

The currently circulating SARS-CoV-2 Omicron variant evades neutralizing antibodies elicited by COVID-19 vaccines.^1,2^ A previous study indicated that BNT162b2 messenger RNA (mRNA) vaccine induces potent cross-clade pan-sarbecovirus neutralizing antibodies in survivors of the infection by SARS-CoV-1, a coronavirus that caused a global SARS outbreak in 2003.^3,4^ However, the ability of these cross-reactive antibodies boosted in SARS-CoV-1 survivors to neutralize the Omicron variant is still unclear.

To address this question, sera samples were obtained from two panels of participants (Fig. S1 and Table S1). The SARS-CoV-1 survivors panel comprised 18 participants with SARS-CoV-1 infection history in 2003. The healthy controls panel contained 21 healthcare professionals from a previously described cohort.^5^ Both panels had received 3-dose of vaccination (two priming doses of CoronaVac followed with one booster dose of protein subunit vaccine ZF2001). For all sera samples, a VSV-based pseudovirus system was utilized to determine neutralizing antibodies against three SARS-CoV-2 variants including prototype virus (D614G), Delta, Omicron (Fig. S2), and SARS-CoV-1 (Tor2), and SARS-like bat coronaviruses WIV-1, on day 0, 14, 90 after third dose vaccination.

The third dose of ZF2001 vaccine rapidly induced a significant increase in humoral immune response. As shown in figure 1A and S3, the geometric mean titers (GMTs) of neutralizing antibodies against the three SARS-CoV-2 variants were significantly increased on day 14 after administration of the third dose in both SARS-CoV-1 survivors panel and healthy controls panel, in consistence with recent third dose booster studies.^5^ A boosting of anti-SARS-CoV-1 and anti-WIV-1 neutralizing antibodies were also observed in SARS-CoV-1 survivors but not in healthy controls (Fig. S3). Importantly, Omicron neutralization titer was dramatically lower than that to D614G (Fig. 1B and C and S4). At 90 days post the third dose, only a half or less were positive for neutralizing antibodies against the Omicron variant in both of the two panels (Fig. 1B). Our results collectively indicate that a 3-dose vaccination is effective at inducing neutralizing immunity to SARS-CoV-2 prototypical D614G and Delta variants but not to Omicron variant even in SARS-CoV-1 survivors tested.

**Figure 1.**
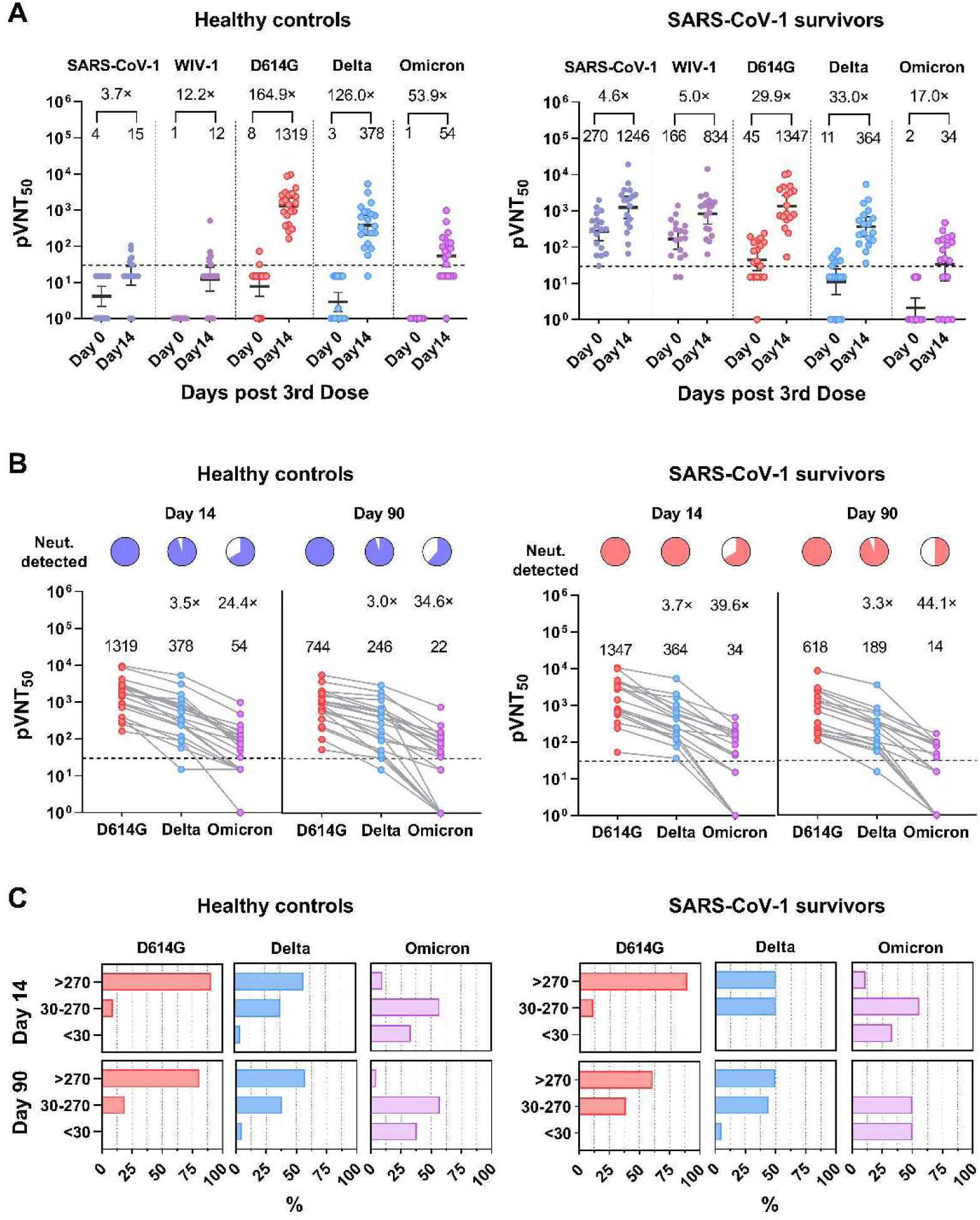
Boosting of Cross-Clade Pan-Sarbecovirus Neutralizing Antibodies but not for Omicron in SARS-CoV-1 survivors. **(A)** Results of pseudovirus neutralization assays using participants’ sera against SARS-CoV-1, SARS-like CoV WIV-1, SARS-CoV-2 D614G strain, Delta strain, and Omicron strain the day before and 14 days after the third vaccination. Neutralizing antibody titers are expressed as sera fold-dilution required to achieve 50% pseudovirus neutralization (pVNT_50_). Dots indicate individual sera samples, dark horizontal lines for each group denote geometric mean titers (GMTs), the error bars indicate the 95% confidence intervals (CI), and the dashed lines indicate the lower limit of detection (LOD, 30). **(B)** Pie charts show the proportion of vaccinees within each group that had detectable neutralization against the indicated SARS-CoV-2 pseudovirus at 14 and 90 days following the third dose of vaccination. Fold-decrease in GMT of Delta and Omicron relative to wild type within healthy controls and SARS-CoV-1 survivors (shown as a number with the “×” symbol); The statistical significance is analyzed by the two-tailed Wilcoxon matched-pairs signed-rank test. pVNT_50_ below the quantitative range but still within the qualitative range (i.e., partial inhibition is observed but a dose-response curve cannot be fit because it does not sufficiently span the pVNT50) was counted half (15) of LOD and no inhibition at all was counted as 1 in statistical analysis. **(C)** The percentages of pVNT_50_ in bar plots after stratification in low (<30), medium (30–270), or high (>270) neutralizing antibody titers are shown for D614G, Delta, and Omicron in healthy controls and SARS-CoV-1 survivors panels.

Potent pan-sarbecovirus neutralizing antibodies are elicited in survivors of SARS-CoV-1 infection after the BNT162b2 mRNA vaccination.^4^ In our research, we also found a relatively broad spectrum of neutralizing antibodies boosted by a third dose vaccination in SARS-CoV-1 survivors. However, these antibodies exhibited dramatically reduced neutralization to SARS-CoV-2 Omicron variant, which indicates that the development of pan variants-adapted vaccines is warranted.

**Figure S1.**
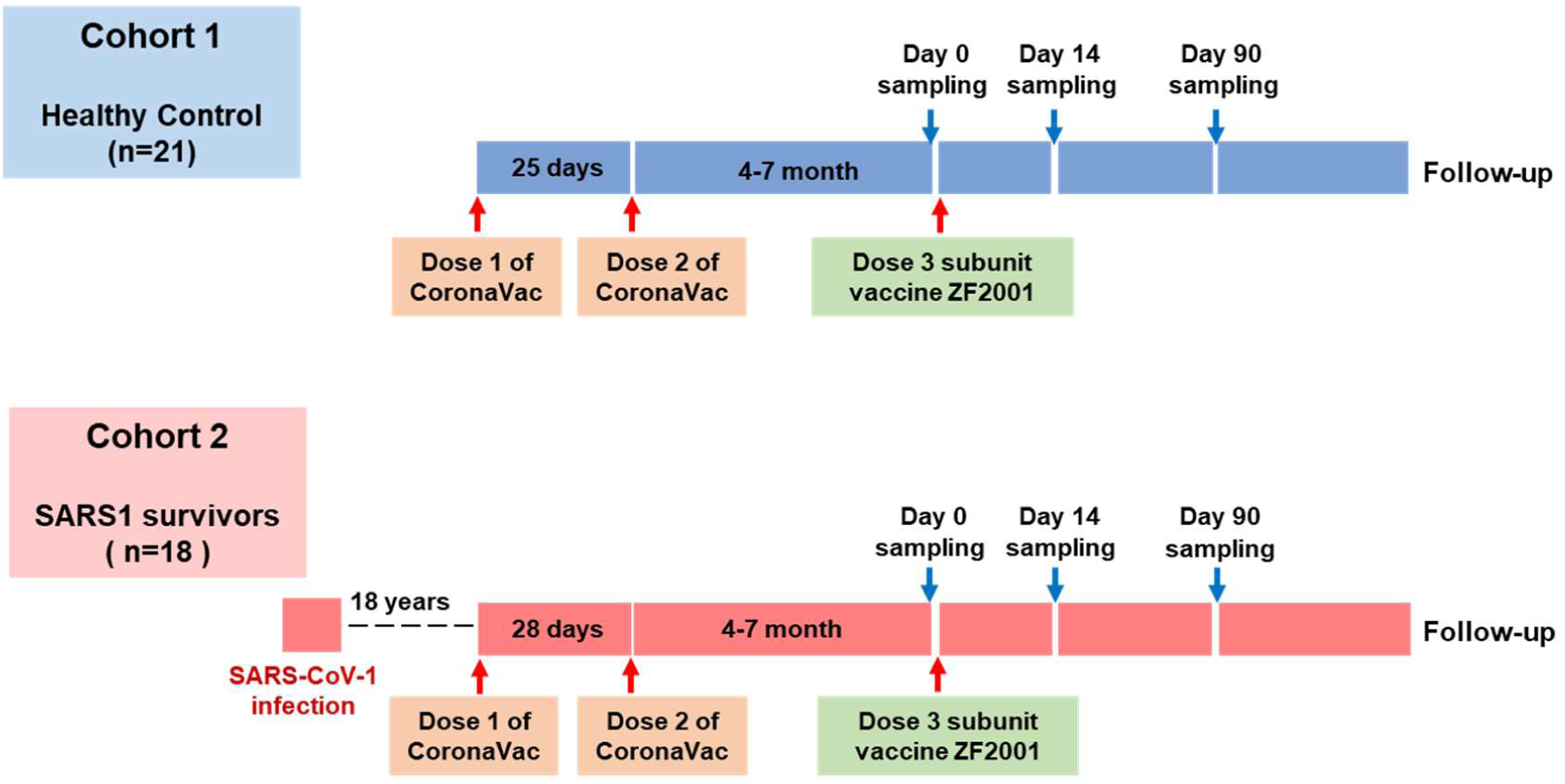
Schematic of trial process timeline. Sera samples were obtained from two panels of participants. The healthy controls panel comprised 21 healthcare professionals at Beijing Ditan Hospital from a previously described clinical trial cohort^5^ (healthy controls). The SARS-CoV-1 survivors panel comprised 18 participants who had been infected with SARS-CoV-1 18 years ago (SARS-CoV-1 survivors). Both of the two panels had received 3-dose of vaccination (two priming doses of CoronaVac in a 28-day interval 4–8 months earlier before one booster dose of protein subunit vaccine ZF2001). Sera samples were collected on day 0, 14, 90 after third dose vaccination.

**Figure S2.**
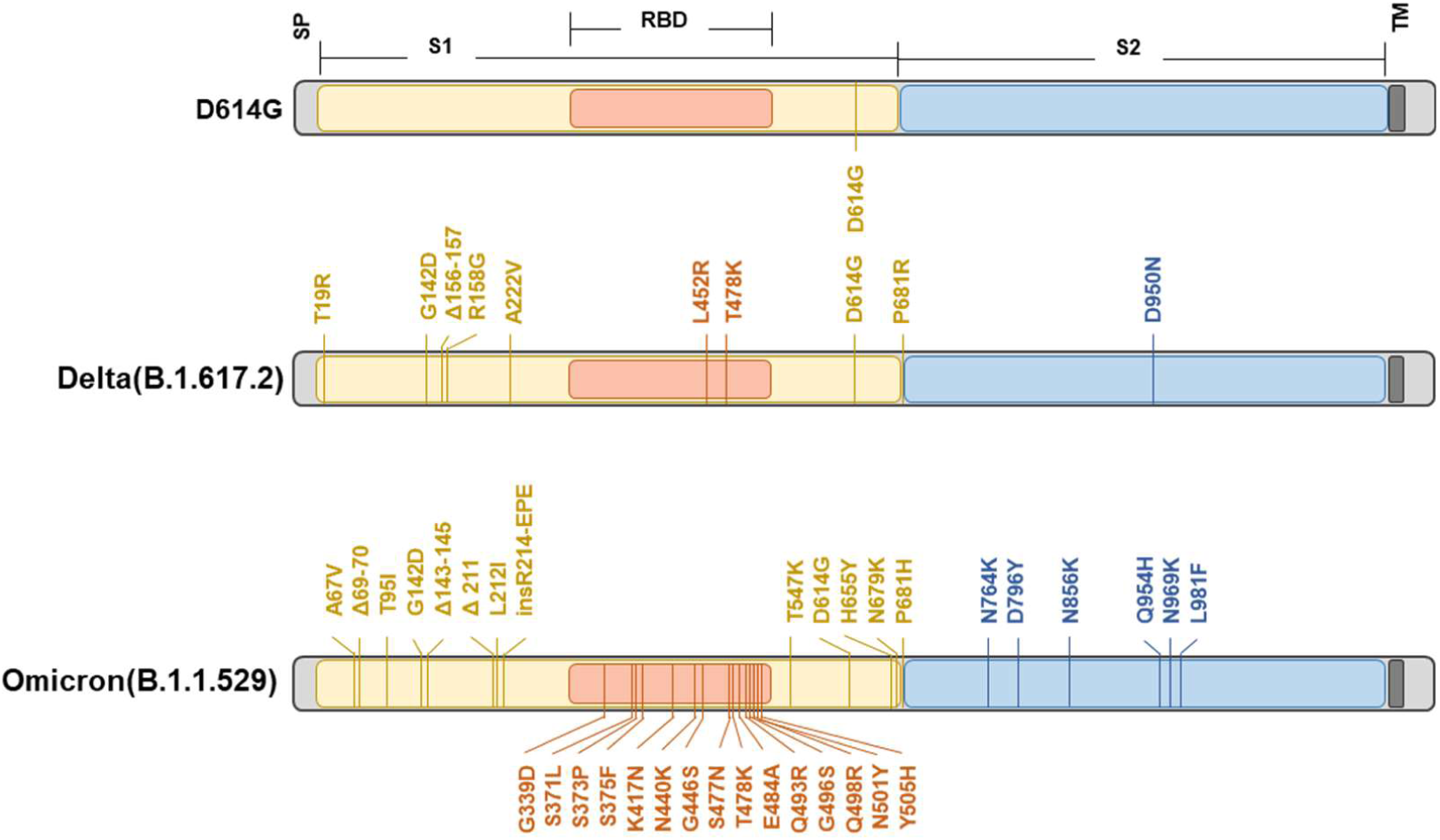
Schematic illustration of the mutations on VOCs spike. Schematic of SARS-CoV-2 spike protein structure and mutations of variants used in this study are illustrated. Omicron variant mutations used in this study were based on the most prevalent mutations (>85% frequency) found in GISAID and reflect the dominant Omicron variant. The regions within the spike protein are abbreviated as follows: SP, signal peptide; RBD, receptor-binding domain; TM, transmembrane domain.

**Figure S3.**
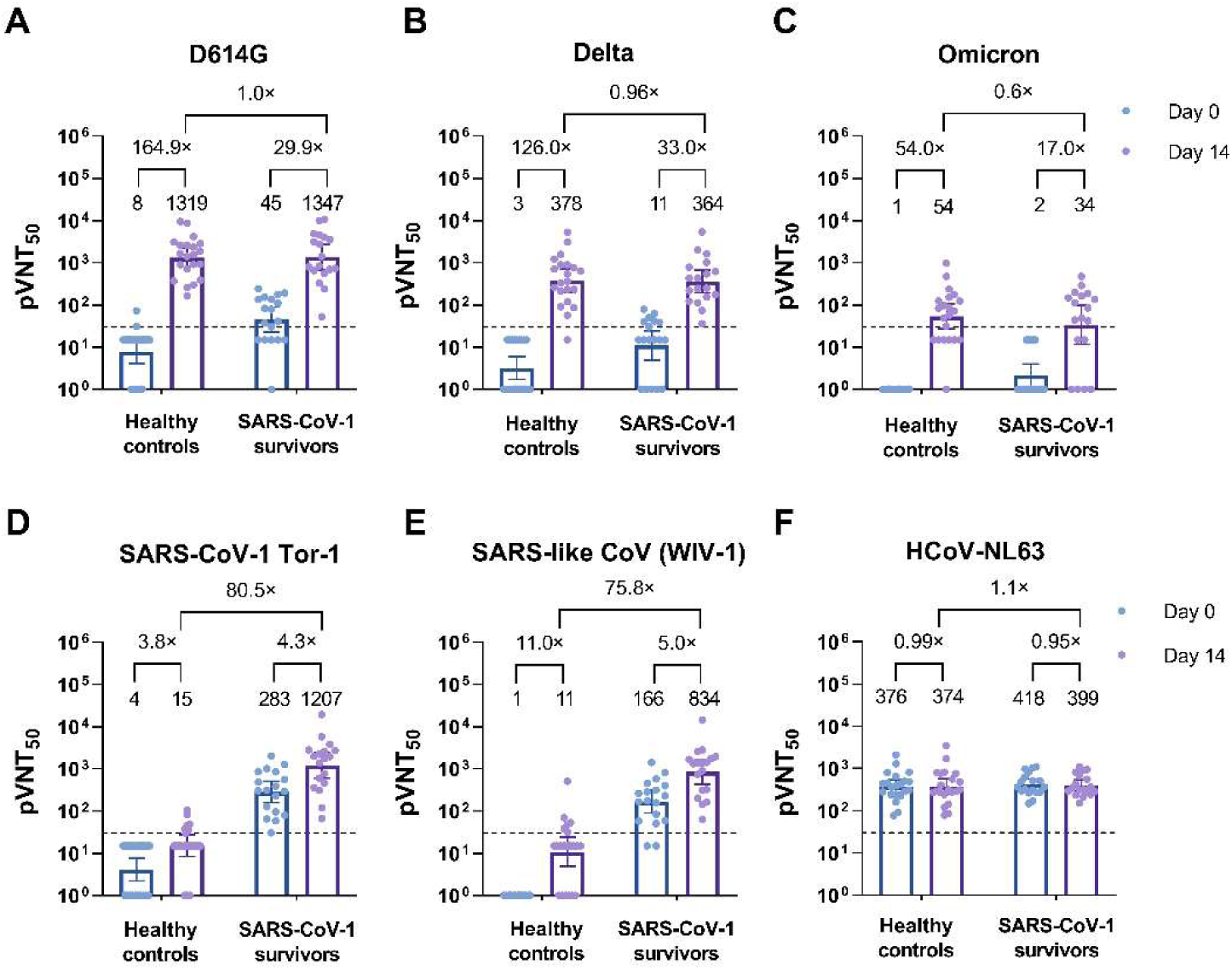
Neutralizing antibody titers (pVNT_50_) against three SARS-CoV-2 VOCs from study participants immediately before (Day 0) or after (Day 14) the third vaccination. Results of pseudovirus neutralization assays using participants’ sera against SARS-CoV-2 prototypical D614G variant (A), Delta strain (B), Omicron strain (C), SARS-CoV-1 (D), SARS-like CoV WIV-1 (E) and HCoV-NL63 (F). GMTs are shown above the bars, and the error bars indicate the 95% CI, and the dashed lines indicate the LOD (30). Fold-increase in GMTs of boosted versus non-boosted individuals is shown as a number with ‘‘×’’ symbol.

**Figure S4.**
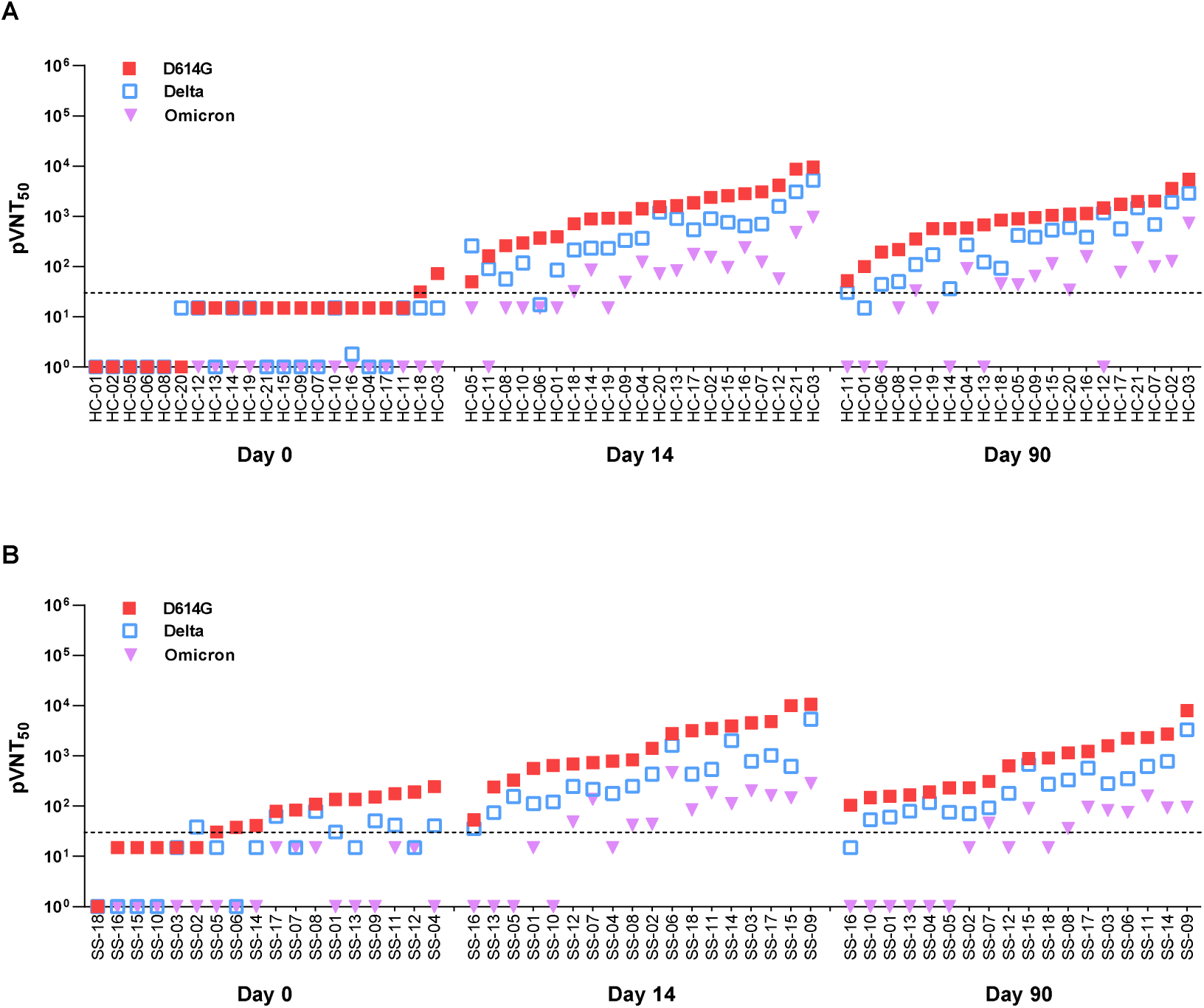
Neutralizing titers against SARS-CoV-2 variants D614G, Delta and Omicron in SARS-CoV-1 survivors and Healthy controls individuals. pVNT_50_ values of sera collected at the day before the third dose vaccination (Day 0), 14 days post the third dose (Day 14) and 90 days post the third dose (Day 90) in Healthy controls panel (A) and SARS-CoV-1 survivors panel (B). Each pVNT_50_ was determined in two independent experiments (each with two technical replicates). The median of the two independent determinations is plotted. Dashed line indicates the LOD.

**Figure S5.**
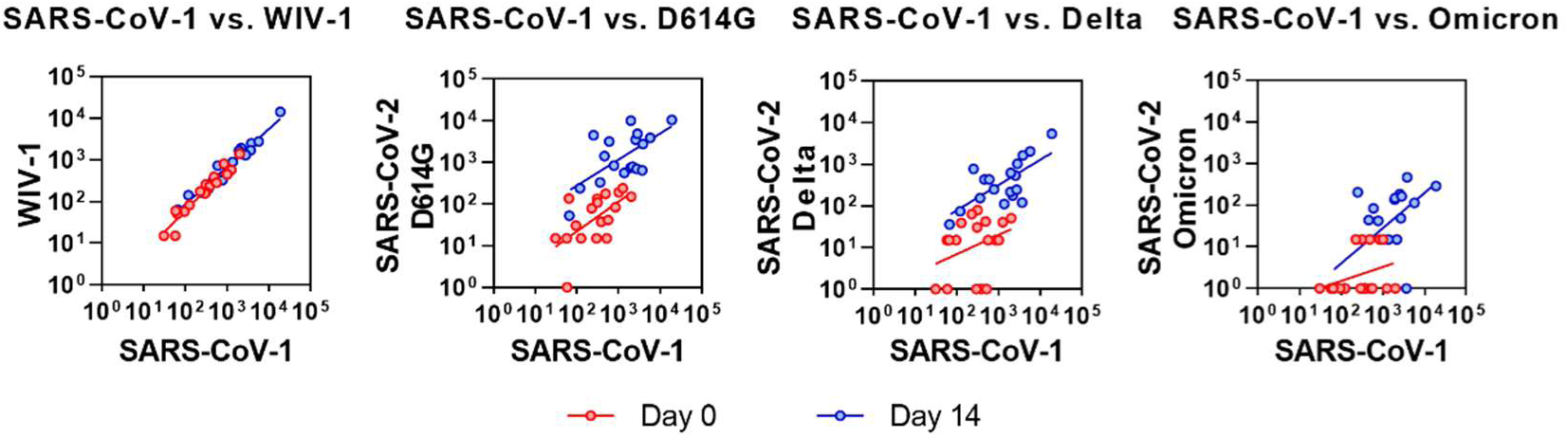
A cross-reactivity of neutralizing antibody response is increased by the third-dose of vaccination in SARS-CoV-1 survivors. pVNT_50_ data from SARS-CoV-1 survivors panel participants that received two-dose vaccination series (Day 0; red circles) or were boosted with the third-dose of ZF2001 vaccination (Day 14; blue circles) were used for linear regression analysis of SARS-CoV-1 versus WIV-1 or SARS-CoV-1 versus SARS-CoV-2 VOCs pseudovirus neutralization. SARS-CoV-1 neutralization titers strongly correlated with WIV-1 neutralization at the day before the 3^rd^ dose (R^2^ = 0.9334; slope = 1.022; p < 0.0001) and at 14 days post the 3^rd^ dose vaccination (R^2^ = 0.9519; slope = 0.9022; p < 0.0001). SARS-CoV-1 neutralization titers correlated with D614G neutralization at the day before the 3^rd^ dose (R^2^ = 0.3742; slope = 0.7148; p < 0.01) and at 14 days post the 3^rd^ dose vaccination (R^2^ = 0.4132; slope = 0.6237; p < 0.01). SARS-CoV-1 neutralization titers showed no significant relationship with Delta neutralization the day before the third-dose of vaccination (R^2^ = 0.1131; slope = 0.4584; p = 0.17); however, ‘‘ZF2001-boosted’’ individuals showed a significant correlation with Delta neutralization titers (R^2^ = 0.4739; slope = 0.6053; p < 0.01); SARS-CoV-1 neutralization titers showed no significant relationship with Omicron neutralization the day before the third-dose of vaccination (R^2^ = 0.1050; slope = 0.3396; p = 0.19) and showed a little bit of correlation at 14 days after the third-dose of vaccination (R^2^ = 0.3199; slope = 0.8464; p < 0.05)

**Table S1.** Demographics of study participants included in this study.

| Panel | Healthy controls | SARS-CoV-1 vaccination |
| --- | --- | --- |
| Number of participants | 21 | 18 |
| Age (Median, range) | 39 (22-57) | 62(39-76) |
| Males | 2 | 8 |
| Females | 19 | 10 |
| Days interval between the second and third doses (Days, Median, range) | 185 (138-226) | 144 (42-228) |

## Materials and Methods

### Ethical statement

The study protocol was approved by the Ethics Committee of the Institute of Beijing Ditan Hospital, Capital Medical University (IRB#2021-(024)-02). Written informed consent was obtained from all participants before the enrollment. This trial was registered with ChiCTR2100051998.

### Sera samples

The sera samples of SARS-CoV-1 convalescents and healthy healthcare workers were provided by Beijing Ditan Hospital, Beijing, China. Sera samples were classified into two panels (Fig. S1): **1)** SARS-CoV-1 survivors received three doses of heterologous vaccines (two priming doses of CoronaVac in a 28-day interval 4–8 months earlier before one booster dose of a protein subunit vaccine ZF2001); **2)** Healthy medical workers received three doses of heterologous vaccines with the same immunization strategy. Participants were sampled at the day before the 3^rd^ vaccination and invited for follow up visits at approximately 14 and 90 days. Detailed information is available in Table S1.

### Cell transfection and pseudotyped virus production

The pSectag2 vector was used to construct recombinant plasmid of codon optimized spike proteins of SARS-CoV-2 prototype (Wuhan-1 reference strain containing D614G mutation) and variants, with a 19 amino acid truncation at the C-terminus of the spike protein^1^ (with mutations shown in Fig. S2). HEK-293T cells were transfected with the plasmids expressing different S protein respectively. VSV-ΔG-G*-Luc pseudovirus *(Kerafast, Boston, MA)* was added 6 h after the transfection. 24 h after the transfection, the supernatant was replaced with fresh complete DMEM medium. Supernatants were collected at 48 h and 72 h after the transfection, passed through a 0.45 μm filter, aliquoted and stored at -80 °C.

### Pseudotyped virus titration

For titration, TREx-293/hACE2 cells were seeded into 96 well plate with 2 µg/ml Tet (∼2×10^4^ cells per well), incubated at 37°C and discarded the supernatant before use. Next day, pseudoviruses stock were taken out the from −80 °C. Pseudoviruses were diluted starting with a 10-fold dilution in a new 96 well plate, followed by eight 3-fold serial dilutions, and each dilution were made in six replicate wells. The diluents were added to the 96 well plate prepared the day before (100 µl per well) and another 6 wells were set as blank control without virus. After incubation for 18h, the luciferase substrate was added for chemiluminescence detection. The 50% tissue culture infectious dose (TCID_50_) of the pseudovirus is calculated by the Reed-Muench method, the cut-off value is 10 times the value of blank control.

### Neutralization assay

The neutralization assay was performed as previously described^2, 3^. Briefly, sera samples which inactivated at 56°C for 30 min were 3-fold serial diluted commencing with a 30-fold dilution, and each dilution was made in two replicate wells. Virus control wells with only virus and cells were set up in each plate. Equivalent pseudovirus (1300 TCID_50_/ml) was incubated with the sera at 37 °C for 1.5 h, and the mixture was then added into the 96 well plate with TREx-293/hACE2 cells. After 18 h the neutralization assay was developed with a luciferase assay system (Promega), and the relative light units (RLU) were read on a Promega GloMax Luminometer. The neutralization rate (%) was calculated as following:

Neutralization Rate 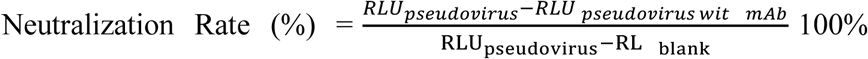. Neutralizing antibody titers are expressed as sera fold-dilution required to achieve 50% pseudovirus neutralization (pVNT_50_). pVNT_50_ was interpolated from the neutralization curves determined using the log(inhibitor) vs. normalized response -- Variable slope fit using automatic outlier detection in GraphPad Prism Software.

### SARS-CoV-2 Spike Variants

This study utilized three SARS-CoV-2 spike variants. The D614G (B.1) variant contained D614G as the only spike mutation. The Delta (B.1.617.2) variant contained spike mutations T19R, G142D, Δ156-157, R158G, A222V, L452R, T478K, D614G, P681R, D950N. The Omicron (B.1.1.529) variant contained spike mutations A67V, Δ69-70, T95I, G142D, Δ143-145, Δ211, L212I, +214EPE, G339D, S371L, S373P, S375F, K417N, N440K, G446S, S477N, T478K, E484A, Q493R, G496S, Q498R, N501Y, Y505H, T547K, D614G, H655Y, N679K, P681H, N764K, D796Y, N856K, Q954H, N969K, L981F. The spike mutations listed here were present in corresponding pseudovirus used in this study.

### Data and statistical analyses

Geometric Mean Titers (GMTs) with confidence interval (CI) of 95% were performed using GraphPad Prism 9.0. pVNT_50_ below the lower limit of detection (<30) was recorded as 15 and no inhibition at all was counted as 1 in the geometric mean calculation. Wilcoxon matched-pairs signed-rank test was performed to detect significant differences in neutralizing titers between the prototype containing D614G mutation and the other variants as well as in titers before and after the third dose. The Bonferroni correction was applied to correct for the increase in type 1 error from multiple testing (adjustment for multiplicity).

## Acknowledgments

This work was supported by the funding from Beijing Municipal Science & Technology Commission, (No. Z201100007920017) to R.J. and National Science Foundation of China (81772173 and 81971916) and National Science and Technology Major Project of China (2018ZX10301-408-002) to X.Z.,National Science Foundation of China (82102363) to D.C. and Beijing Natural Science Foundation (M21007) to J.D.

## Authors’ contributions

X.Z., X.W., and R.J. conceived, designed and supervised the experiments; X.Z., D.C. and X.W. wrote the manuscript; D.C., Y.Q., X.L., and Y.S. performed the neutralization experiments. R.S., X.H., J.D., Y.Z., and F.X. provided convalescent sera and patients information. All of authors approved the final manuscript.

